# General Theory of Metabolism related to Animal’s Taxonomy and Size

**DOI:** 10.1101/2025.04.08.647784

**Authors:** Peter F. Pelz

## Abstract

Each mammal has a budget of approximately one billion heartbeats after birth. This is consistent with their heart rate and life span, which scale with their mass to the power −1/4 and 1/4 respectively, given a complex cardiovascular system. However, the underlying empirical law, i.e., Kleiber’s law, according to which the metabolic rate scales with the mass to the power of 3/4, applies to all animals: for instance, flatworms with a most simple vascular system and a size slightly above the diffusion limit on growth show the same metabolic scaling as mammals. To date, there is no concise theory that is consistent with cell metabolism and compatible with physiological laws, e.g., that the volume flow scales with the capillary diameter to the power of three (Murray’s law). In this paper we present how cell metabolism determines the scaling of the organism’s metabolic rate via the *Metabolic Module* (MM), a cylinder formed by the organism’s cells with a concentric capillary – sized and shaped with scarcity. Evolutionary changes from one taxonomic class to the next led to an unsteady increase in the number of MMs: the metabolism of protists and planarians, e.g., flatworms, is given by one MM only; for ectotherms, i.e., cold-blooded organisms, one thousand and for endotherms, i.e., warm-blooded organisms, nearly one hundred million MMs working together. Special cases, such as diffusion-limited metabolism and the 3/4 power law are asymptotes of the presented general theory. The presented general theory of metabolism offers valuable insights for the targeted development of artificial tissues.

## 1 Introduction

In the early 1930s, the Swiss biologist Max Kleiber expected a slope of approx. 2/3 when he plotted the measured basal metabolic rate *Ė* of mammals against their body mass *m* in a log-log plot [1] (Fig. 3). He suspected a size constraint due to Galileo’s “*surface law*” [2] in connection with heat transfer. This would imply, for a given metabolic rate, the surface area limits the volume or mass *m* of the organism. For geometric reasons, the surface area is proportional to body volume to the power 2/3. The expected relationship was therefore *Ė* ∝ *m*^0.66^. However, he noted: “*The best-fitting unit of body size for comparing the metabolism of rat, man, and steer has been found to be m*^3*/*4^.” [1]. Today, the empirical relationship *Ė* ∝ *m*^3*/*4^ (Eq. 9) is known as Kleiber’s law [3]. This relationship is visible in the measurement data for large body masses (symbols in Fig. 3) and in the asymptotes (broken line labelled Eq. (9) in Fig. 3) of the presented absolute, i.e., not relative, theory (solid lines in Fig. 3).

To date, there is no (i) complete theory of metabolism related to mass and taxonomy that covers the range from one picogram to 100 tons; there is no (ii) concise theory based on only a small number of assumptions; and there is no (iii) consistent theory based on cell metabolism and the physicochemical properties of cell tissue that is also consistent with Murray’s law. This is the Pareto optimal solution to the multi-objective optimisation problem, where mass and dissipation are to be minimised simultaneously at a given flow rate. This empirically validated relation states that the flow rate through a capillary is proportional to its diameter to the power of three [4], [5], [6]. After all, the existing theories are all only relative in the sense of the scaling equation *Ė*_2_*/Ė*_1_ = (*m*_2_*/m*_1_)^0.75^. As a result, their scope is limited to scaling the metabolic rate from one mass *m*_1_ to another mass *m*_2_. In contrast, the theory presented here allows the absolute value of the metabolic rate to be determined as a function of mass, taxonomy and physiological constants: *Ė* = *Ė* (mass, taxonomy, physiology).

To date, there have been three approaches to explaining Kleiber’s law. The first assumes that the metabolic rate for a given mass is limited by heat transport. The second assumes a limitation due to the supply of nutrients or oxygen. The third states that the metabolism of the cell itself depends on the size of the organism [7]. Only the last approach establishes a link to cell metabolism, but I doubt that this provides an explanation for the entire mass range from one picogram to 100 tonnes.

As for the first approach, Max Kleiber was wrong with his hypothesis that the heat transport over the body surface limits the metabolic rate. Measurements show that the second approach is appropriate: without reasoning to be derived in this paper the empirical data in Figure 5 reveal that the heart rate of mammals scales as *f* ∝ *m*^−0.25^. Since the relative heart volume and the total number of heartbeats are dimensionless, both variables must be scale-invariant for reasons of dimension. This follows from the Bridgman postulate [8], which demands the “*absolute significance of relative magnitude*”. Both statements are confirmed by empirical data: heart volume accounts for about 6% of body volume in all mammals [9], and the number of heartbeats is about one billion [10] (Fig. 5). Since the blood volume flow is given by the product of heart volume and heart rate, it scales as *m*^0.75^.

Recently, when modelling metabolic rate, a linear combination of the 2/3 power of mass with a linear mass term was discussed [11], which follows the first approach. We interpret this rather as a polynomial interpolation of measured data that lacks a physicochemical basis.

The second approach has been pursued since 1997 with the WBE theory [12] as a starting point: Kleiber’s law is derived from the topology, shape, and size of the distribution network of nutrients and oxygen. The WBE and similar theories [13] assume that the diameter of the capillaries is scale-invariant and that the vascular tree is large and highly branched. Therefore, neither theory can explain the metabolism of very simple organisms such as planarians, which do not have a highly branched vascular system. The WBE theory requires eight assumptions to derive the 3*/*4 power law [14]. It is therefore neither concise nor elegant in the sense of (the philosophy of science known as) Occam’s razor [15], [16], [17]. Finally, all mentioned theories disregard cell metabolism when they assume a constant capillary diameter. In fact, a scale-invariant capillary diameter conflicts with Bridgman’s postulate [8]; and indeed, the measured capillary diameters increase with the size of the organisms [18].

I analytically explain the empirical Kleiber’s law solely by the only demand for efficiency, consistent with cell metabolism and Murray’s law. The *cybernetics of evolution with scarcity* (Fig. 2) is the conceptual framework for the *general theory of metabolism* as a function of size and taxonomy. This framework builds on known cybernetics of evolution [19]. In the extended framework presented here with social, design and quality space (Fig.2), scarcity drives nature’s *evolution for efficiency* and *evolution for innovation*. This results in a Pareto optimally sized and shaped *Metabolic Module* (MM), a cylinder formed by the tissue cells for the organism’s metabolism. Suppose a cell is the smallest scale building block, then the MM is the mesoscale building block to compose an organism by up-numbering the Pareto optimally sized and shaped MM. The modules considered so far are genes, genomes, organelles, cells and organs. The MM is now placed between cells and organs.

The quantitative general theory based on the up-scaled and up-numbered Pareto optimal MM predicts the metabolic rate of organisms with a given mass, confirmed by empirical data ranging from one picogram to 100 tonnes.

The general theory has two asymptotes: the *diffusion asymptote* for ‘small’ and the *convection asymptote* for ‘large’ organisms. For the diffusion asymptote the metabolic rate is proportional to the mass, for the convection asymptote it is proportional to the 3/4 power of the mass. Therefore, Kleiber’s law is a special case of the general theory.

## 2 Overcoming the Diffusion Limit on Growth

Life is about survival and sustaining genes. The discussion on this philosophic hypothesis is clearly beyond the scope of this paper. Regardless of the outcome of this discussion, it is clear that survival and sustaining requires resources for metabolism, and as resources are (usually) scarce, efficient use is essential. In times of scarce resources and given cell metabolism, the locomotion of organisms therefore requires firstly a high propulsion efficiency on land [20], in water [21] or in the air [22] and secondly a minimum body mass for a given power output in order to win the battle for resources or to escape from a resource: the realised function of high acceleration or speed offers advantages in competition for scarce resources. (The objective of minimising mass is also important when considering transformation rather than locomotion. For example, the energy and material resources available for incubating an egg are limited from the very start.)

As mentioned, the cybernetics of evolution with scarcity (Fig. 2) is a conceptual framework. It helps to develop the general theory of metabolism given in the following and it allows the localisation and distinction of strategies such as sufficiency, efficiency, innovation (Fig. 2b), cooperation and competition (Fig. 2a). It enables a distinction to be made between allocation Pareto optimum, Nash equilibrium (Fig. 2a), and quality Pareto optimum (Fig. 2d). The feedback loop separates the life cycle of an individual (Fig. 2c) from the interaction within and between species in the social space (Fig. 2a). The conceptual framework makes clear the difference between a ‘Darwinian’ strategy and a ‘Newtonian’ constraint. Both were falsely discussed as contradictory [23] because a cybernetic picture such as the one presented here was previously lacking.

The following discusses the constraints of the design space and the need to overcome the diffusion constraint on growth. The cell is at the end of the supply chain of metabolism. At the same time, they are the first and basic building blocks of organisms and their physicochemical properties (cell metabolism, diffusive permeability) are similar within and between species.

Physicochemical laws, available modules and materials, allocated and therefore available resources and the required function form the design space (Fig. 2b). Suppose competition in the social space (Fig. 2a) triggers a functional need satisfied by growth. The direction of this process is the opposite of sufficiency. The design space is therefore shrinking due to the need for size. In the limiting case, the space shrinks to a point that represents the *limit on growth* of the organism for given laws, modules and resources. With the given laws and resources, the growth limit can only be overcome by new modules or materials.

What if cells are the only modules available? Suppose *r* = *k*Δ*c* is the consumption of oxygen molecules of concentration Δ*c* per unit of volume on a cell level. In this case, *R* = *rV* molecules per unit of time are needed for the organism’s metabolism, which has a volume size of *V*. The reaction rate *k* gives the consumption speed, and the inverse of *k* is the consumption time. To prevent death by starving, the consumption time must be larger than the supply time. From that aspect, the most vulnerable cells have the largest distance *δ* from the source of metabolism, the cell’s interface to the ambient.

Transport within the cell tissue is diffusive. Hence, the order of magnitude of the time needed for transporting metabolism to the cells is *δ*^2^*/*𝒟, were 𝒟 is the diffusion coefficient in Fick’s law. Only provided *δ*^2^*/*𝒟 *<* 1*/k*, is diffusion fast enough to prevent of the most distant cells starving. The diffusion constraint on growth 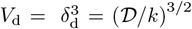 is reached when the diffusion time equals the consumption time. For a ‘small’ organism, i.e., for an organism with volume size *V ≪ V*_d_, the kinematic metabolism rate *Q* = *R/*Δ*c* is proportional to the size *Q* ∝ *V*:

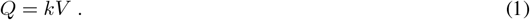

The dimension of *Q* is volume per unit of time. For the general theory of metabolism developed presented in this work, *Q* is also a volume flow. With the energy density *e* = |Δ*G*^*°*^| Δ*c*, the molar Gibbs free energy |Δ*G*^*°*^| of the cell metabolism, and the mass density *ϱ* = *m/V* of the cell tissue, Equation (1) is written as energetic metabolic rate *Ė* = *ke m/ϱ*. We call this well-known equation the *diffusion asymptote*. The metabolic rate of ‘small’ organisms, i.e., protists like the plankton shown in Figure 1a, is given by this diffusion asymptote as revealed by Figure 3.

**Figure 1:**
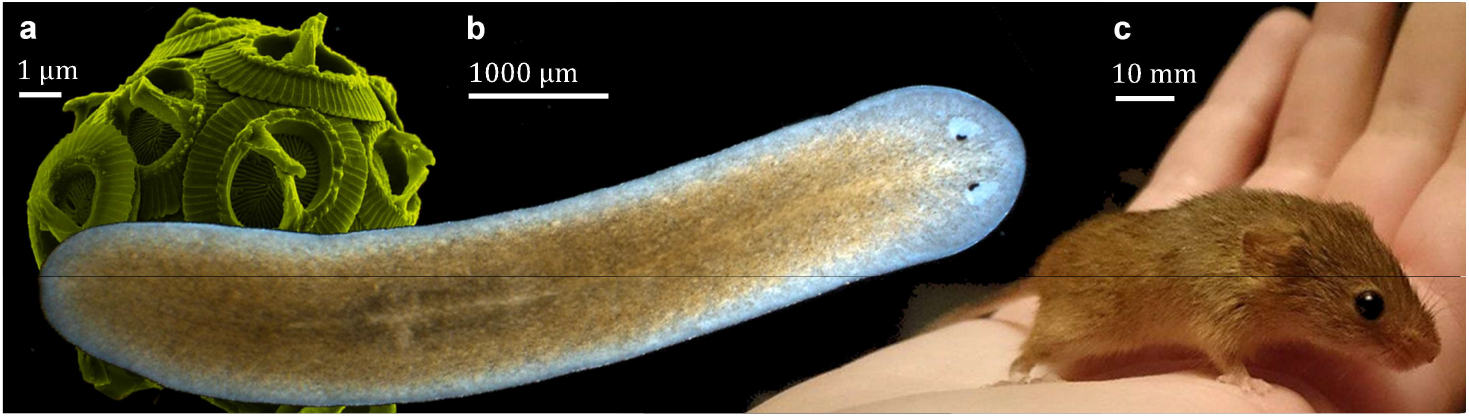
Organisms of different taxonomy, size, and total numbers of Metabolic Modules (MMs) **(a)** Protists represented by a self-feeding single-celled organism of the plankton community (photo taken from https://commons.wikimedia.org/wiki/File:Gephyrocapsa_oceanica_color.jpg); **(b)** a flatworm or planarian with one MM only formed by the gastrovascular cavity (photo by and with permission of Jochen Rink [7]); **(c)** during evolution the MMs are ‘up-numbered’ by nature with up to one billion MMs now part of the vascular system operating in parallel. In any case, the MM is Pareto optimal, i.e., size and shape evolved such that the resources are optimally utilized (photo by and with permission of Fernanda Ruiz Fadel).

**Figure 2:**
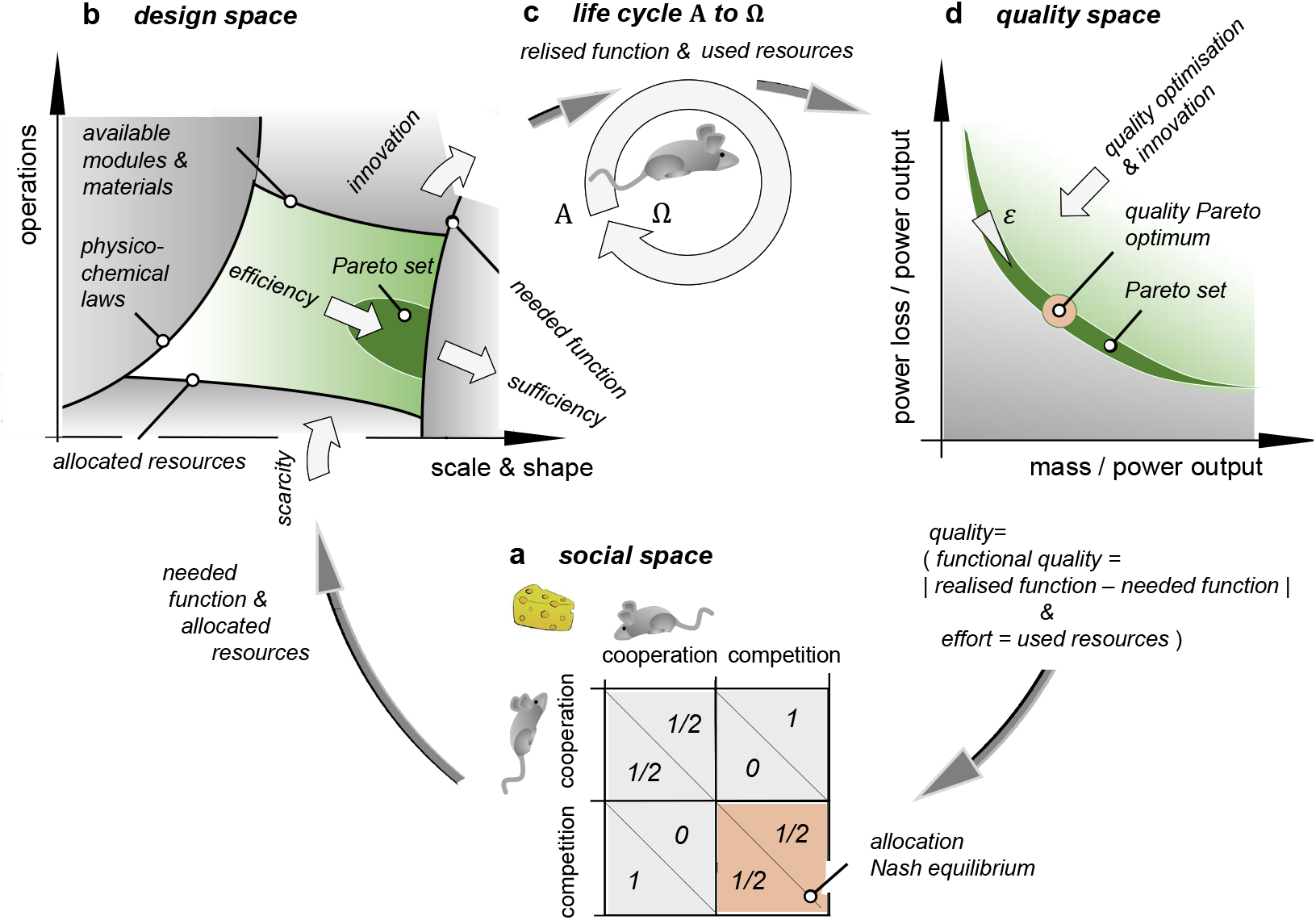
Cybernetics of Evolution with Scarcity. The system of interest is the organism’s life cycle **(c)**. The controller has the social space **(a)** and the design space **(b)** as serial processes. A point in the design space - given by operating and design parameters - is mapped via the organism’s life cycle **(c)** to a point in the quality space **(d)** closing the loop. Feedback loops of organisms of the same or different species intersect in the social space **(a)** where cooperation or competition can result in the allocation Nash equilibrium or allocation Pareto optimum (not shown). This allocation optimum or allocation equilibrium of different species differ from the quality Pareto optimum of one species. The latter is optimal concerning functional quality, i.e., the difference between realised and needed function, the robustness of the functional quality, and the effort measured in used resources.

**Figure 3:**
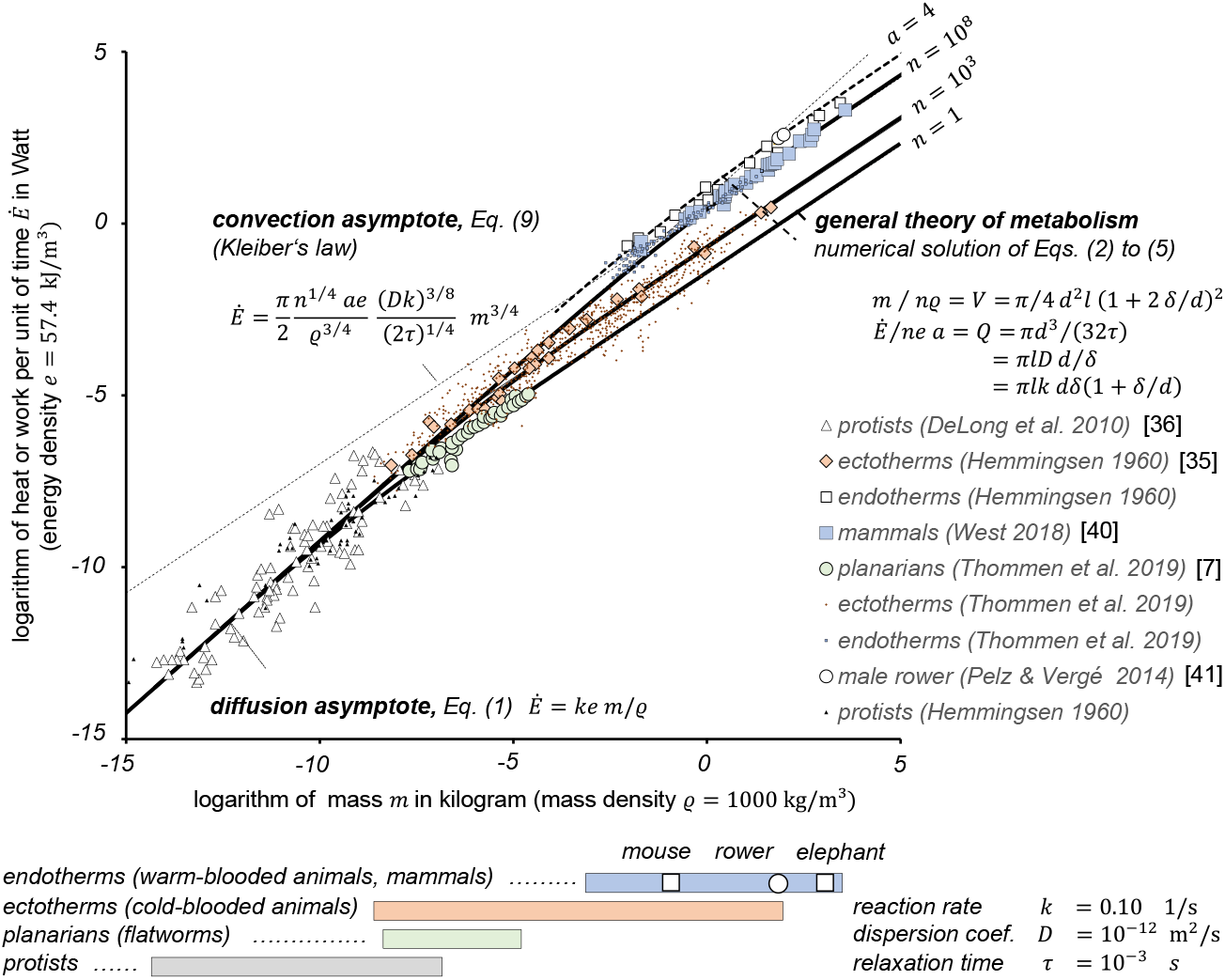
General Theory of Metabolism and their asymptotes - predicted and measured metabolic rate vs. body mass; Metabolism is measured in work or heat per unit of time vs. body mass. The symbols show the measured metabolism of protists, planarians, or flatworms, ectotherms or cold-blooded animals, the basal metabolic rate of endotherms or warm-blooded animals, and the active metabolic rate of mammals including rowers. The thick solid lines show the Pareto optimal metabolic rate, i.e., the solution of Eqs. (2) to (6). The thin broken lines show the diffusion asymptote and the convection asymptote given by Eq. (1) and Eq. (10) respectively for the basal metabolic rate. The dashed line is the Pareto optimal metabolic rate for mammals with an activation factor of four. All other lines show basal metabolic rate with an activation factor of one.

If an organism tends to grow beyond its limit on growth, *V > V*_d_, diffusion alone is insufficient to prevent starvation. Now, the danger of death is the strongest innovation pressure one can imagine. To release that innovation pressure, nature invented *“blood [that] is a juice of exceptional kind”* as Mephistopheles stated in Goethe’s Faust [24].

In more general terms - independent of blood as a fluid - the innovation compared to purely diffusive mass transport is convective mass transport. The supply chain becomes sufficient to prevent starvation if the short-distance diffusive transport along the ‘last mile’ is downstream of the long-distance convective transport by blood cells. Blood cells store and transport oxygen like lorries carrying mail to the distribution points. From there, postmen deliver the mail on the ‘last mile’, analogous to the diffusion process through the tissue.

The innovation pressure to overcome the diffusion constraint on growth is explained with the cybernetics of evolution with scarcity (Fig. 2). Size, shape, and operations define the dimensions of the unconstrained design space as sketched in Figure 2b. The accessible design space with all feasible combinations of size, shape, and operating conditions covers only a small room, the design space as such. The design space has four walls or constraints. These are given by (i) the allocated and thus available resources, (ii) by the physicochemical laws, (iii) by the needed function such as speed to escape enemies or chase victims, and (iv) by the available building blocks, i.e., the available modules and materials.

As discussed above, resources are allocated in cooperation or competition with other species of the same or different kind, usually in a mathematical game setup. Functions like speed to escape enemies, hunt victims, or attract a partner are needed to win this ‘*game of life*’ in the social space (Fig. 2a). Therefore, the two constraints (i) available resources and (iii) required function are at least partially interdependent due to the feedback loop.

Reducing the functional need in a freely made decision is called sufficiency. Sufficiency widens the design space as sketched in Figure 2b and allows for solutions with lower resource consumption. Down-steaming, i.e., speed reduction is a typical example of sufficiency.

Physicochemical laws such as Lavoisier’s law of conservation of mass, Carnot’s second law of thermodynamics, Meyer’s and Joule’s first law of thermodynamics, Newton’s and Euler’s laws of momentum, and the associated constitutive relationships such as Fick’s law of molecular diffusion naturally constrain the design space.

Before reducing needs by employing sufficiency, the efficiency and innovation pressure gives two strategies usually applied first: (i) evolution for efficiency is a heuristic search within the given design space for optima that takes many overall feedback loops. In contrast, (ii) evolution for innovation widens the design space (Fig. 2b). It ‘pushes’ the limit on growth, i.e., the above-mentioned point in the design space, through newly available modules or materials (while at the same time complying with physicochemical laws). In the evolutionary process, innovation is therefore a preferred alternative to sufficiency in order to fulfil needs with resources through functions. Since efficiency comes first, the order of priority is to explore the design space (efficiency), before expanding the design space with new modules or materials (innovation), before expanding the design space by reducing needs (sufficiency).

In the light of the presented cybernetics of evolution, Darwin’s evolution is a strategy in the design space (and social space), and axioms such as Newton’s second and third law are constraints on the design space. In other words, Darwin’s name stands for evolution for efficiency or innovation. Both are heuristic techniques. Newton’s name stands for physicochemical constraints, i.e., axioms. Therefore, there is no conflict between a ‘Darwinian’ and a ‘Newtonian’ approach [23], but rather an interaction in the cybernetics of evolution.

In fact, at the heart of this paper is the understanding of the mesoscale but scale-variant building block, the MM. This module, expands the design space and enables the diffusion limit on growth to be overcome. The MM forms the basis for the general theory developed in the section after next. First, however, the existing metabolic scaling theory will be evaluated.

## 3 Sizing and Shaping the Metabolic Module with Scarcity

Occam’s Razor demands conciseness and this is not only a model quality, as described above, but also a design quality. Design is a multi-stage, constrained optimisation problem; first, a feasible specific topology is selected; second, the sizing and shaping happen; in the third step, the operating parameters are selected [25]. Conciseness concerns the first step, the decision for one specific topology. The claim *“less but better”* by the influential designer Dieter Rams [26], just as the following aphorism of Antoine de Saint-Exupéry [27], is an expression of Occam’s Razor for the design of systems and modules:

> *“Perfection is finally attained not when there is no longer anything to add, but when there is no longer anything to take away*.*”*

Regarding the design of the MM: Figure 4 shows the simplest topology satisfying the need for convective transport with the interchange to diffusive transport. The most likely module is a cylinder of length *l* whose tissue of thickness *δ* encloses a concentric capillary of diameter *d*. A large number *n* of such modules are supplied in parallel by the metabolism. For mammals, the total number of MMs has an order of magnitude of one billion [12].

**Figure 4:**
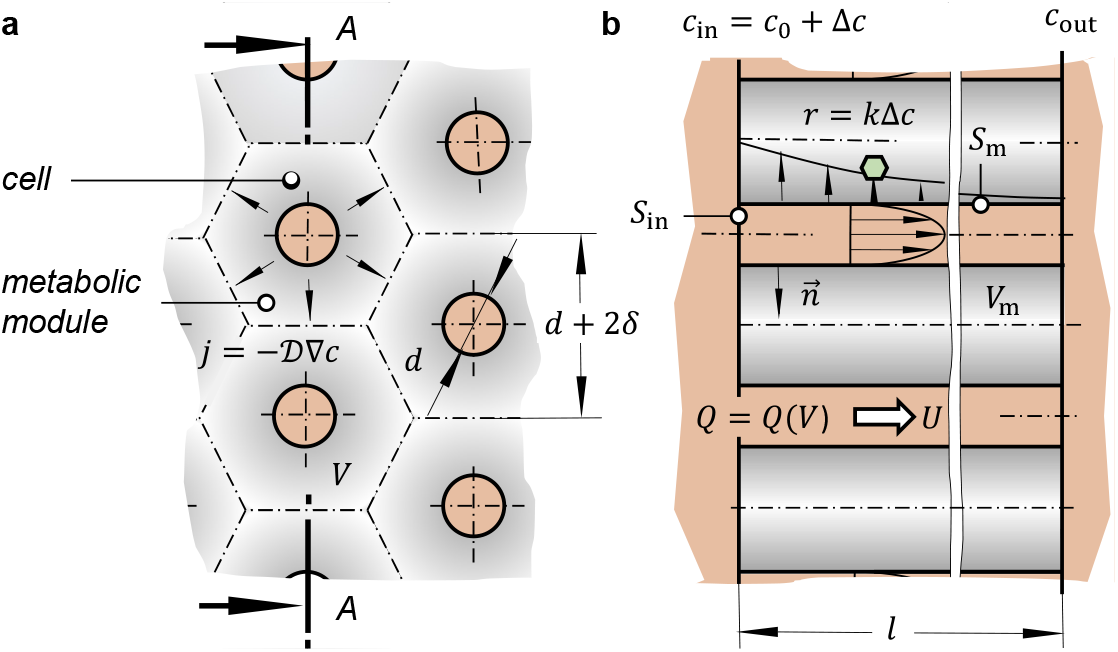
(a) Tissue composed of Metabolic Modules (MMs) arranged and operating in parallel; **(b)** cross sectional view A-A of the MMs

**Figure 5:**
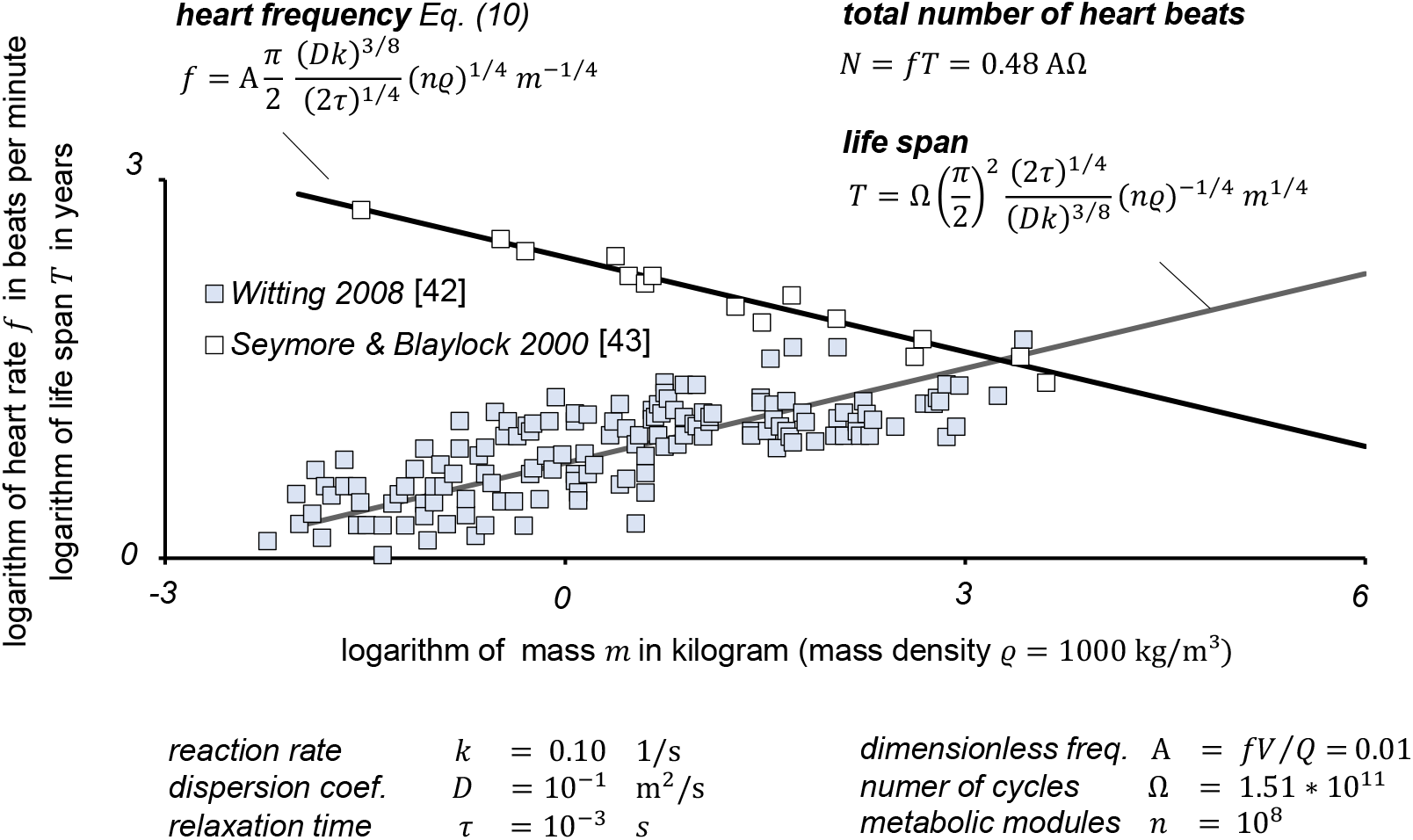
Predicted and measured mammals’ heart rate and life span vs. their body mass.

With respect to Figure 4, the shape parameters are *δ, d* and *l*. The size is given by the volume

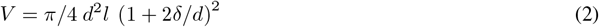

for a circular cylinder a cross-sectional area corresponding, for example, to that of the hexagon cylinder shown in Figure 4. The operating design variable is the blood speed *U* and the resulting volume flow *Q* = *π/*4 *d*^2^ *U*.

Assuming *Q* to be given, we derive the size and shape of the MM for severe scarcity, i.e., a shrinking design space, (Fig. 2b). Besides Equation (2) three more constraints or optima are needed to determine the four unknowns *V, d, l* and *δ*.

First, we have the Pareto optimum of two competing objectives: animal locomotion under scarcity fosters efficient lightweight solutions. The mass *ϱV* and the dissipation due to viscous friction shall be minimal. The flow may pulsate with a heart rate *f*. For the flow within the capillary the Womersley number 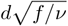 is much small than one for the given kinematic viscosity *ν*. Hence, the dissipation equals the product of pressure loss and volume flow: Δ*pQ* = 8*π νϱU* ^2^*l*. With the relative weight of the two objectives *ε* the resulting constrained multi-objective optimisation problem reads 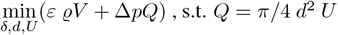.It is sketched schematically in the quality space (Fig. 2d). The problem is equivalent to the unconstrained problem 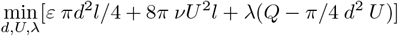 with the Lagrange multiplier *λ*. Using the abbreviation 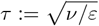,known as Kolmogorov time scale in turbulence [28], the Pareto optimal flow velocity is *U* = *d/*(8*τ*). Multiplying this result by the cross-sectional area *π/*4 *d*^2^ showed that this is in fact Murray’s law, i.e.,

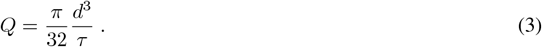

From a mathematical point of view 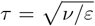is the curve parameter of the Pareto set forming a surface (Fig. 2d). From a physiological point of view, *τ* is a relaxation time for the time-dependent adhesion processes of the blood components at the capillary wall [29], [30]. The reciprocal is the related apparent wall shear rate 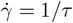.Due to the non-Newtonian nature of blood the real shear rate is slightly higher but of the same order of magnitude. Multiplying the wall shear rate with the blood’s dynamic viscosity *µ* = *ϱν* yields the shear stress at the capillary wall 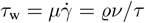.

Nature presumably forms capillaries for a constant wall shear stress by feedback of the latter in the growing phase [29], [30], [31], [32]. Within the capillaries the validated relation *Q* ∝ *d*^3^ holds independently of this mechanism. The general theory of metabolism is built on the Pareto optimum (Eq. 4), that can be considered similar to a physiological constraint. Therefore, all presented results are fully consistent with Murray’s law.

There are three alternative approaches to the following second and third step: We can analyse the coupled convection-diffusion systems for both the fluid, e.g., the blood, and the cell tissue based on its integral formulation. Alternatively, we can solve the two boundary value problems consisting of the differential form of the above equations. This leads to eigenvalue problems and finally to the concentration distribution of, e.g., oxygen in the cell tissue. Finally, the boundary value problem can be solved numerically. In all three approaches, the optimisation is based on the analytical or numerical solution. In principle, there is no difference between the approaches. Due to its conciseness, we solve the reaction-diffusion system in its integral formulation in this paper.

Secondly, for the supply chain, we demand – for an efficient take up – that the residence time of the blood in the capillary is sufficiently long so that the supplied oxygen molecules of concentration *c*_in_ = *c*_0_ + Δ*c* are absorbed by the tissue, i.e., the oxygen concentration at the outlet vanishes. For this first limit, *c*_out_ → 0, the danger of starving cells at the downstream end of the module is severe. If the residence time is small, for the second limit, *c*_out_ → *c*_0_ + Δ*c*, the blood cells would be used insufficiently. The optimal supply is reached if the outlet concentration equals the basal concentration, *c*_out_ = *c*_0_. For this optimum only, Δ*c* is completely consumed and there are no starving cells.

The net convective flux of molecules for this optimal case is (*c*_in_ − *c*_out_)_opt_ *Q* = Δ*c Q*. The principle of mass conservation states that this flux equals the diffusive flux through the surface of the capillary *S*_m_ with surface normal vector 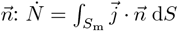 (Fig. 4). With Fick’s law of diffusion relating the molar flux vector to the concentration gradient, 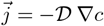,we arrive at 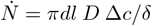.Here we introduce the dispersion coefficient *D* in analogy to G. I. Taylor’s dispersion coefficient for capillary flow [33]. The dispersion coefficient is proportional to the molecular diffusion coefficient *𝒟*, but smaller due to inhomogeneity of the reaction-diffusion effects within the MM. From the balance for the optimal case 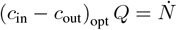 we derive

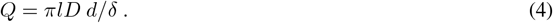

And finally, thirdly, the net and optimal convective molar flux Δ*cQ* transported via diffusion to the cells equals the cell metabolism of all tissue cells: 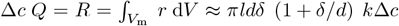. Therefore, the final mass balance reads

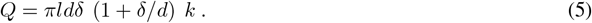

The reaction-diffusion equation for the MM can also be solved numerically and the solution is provided in the supplementary material of this paper. This solution would replace Eqs. (5) and (6). Both models, the presented and the numerical, show the same physicochemical structure and vary only in details.

Eqs. (2) to (5) form a nonlinear system of equations for the four unknowns *V, d, l, δ* that can be easily solved by anyone without any special knowledge of numerical methods simply by using a spreadsheet. For this purpose, *δ/d* is entered as an independent variable in the first column with a logarithmic spacing covering the asymptotic ranges *δ/d≪* 1 and *δ/d ≫* 1. The diameter *d* is derived by dividing Eq. (5) with Eq. (4). The result fills the second column of the spreadsheet. Eq. (3) gives the volume flow *Q*, filling the third column. The capillary length *l* results from Eq. (4) to be plugged into the fourth column. The fifth column is filled with the volume *V* of one module determined from Eq. (2). The Pareto optimal metabolic rate *Q* = *Q*(*V*) is derived by plotting the values of the third and fifth column against each other.

If this is done, two asymptotes of the general solution appear, namely one in which the metabolism is dominated by diffusive transport of the metabolites and one in which it is strongly influenced by convective transport. For *δ/d ≫* 1 the first summand within the brackets of Eqs. (2) and (5) can be neglected yielding the *diffusion asymptote* (Eq. 1). For *δ/d ≪* 1 the second summand can be neglected yielding the *convection asymptote*, known as Kleiber’s law

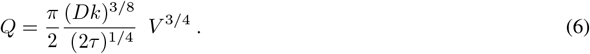

The new physiological constant *π/*2(*Dk*)^3*/*8^ (2*τ*)^−1*/*4^ takes cell metabolism into account. The *convective asymptote* of metabolism is a power law and hence, scale-free in the limit *δ/d≪* 1. The general solution of Eqs. (2) to (5) covers the *diffusion asymptote* and the *convection asymptote*. Therefore, there is a typical volume and a typical volume flow. Two scales are given by the intersection of both asymptotes shown as dotted lines in Figure 3.

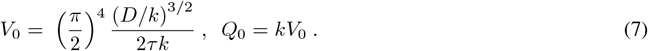

The volume flow *Q* is a kinematic metabolic rate of one MM as a function of the MM’s volume *V* given by Eq. (7). The Pareto optimal MM reveals the frequency scaling

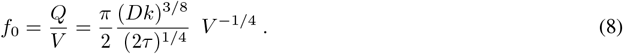

The residence time *l/U* of fluid or blood within the MM is derived independently from Eqs. (2) to (5). For the Pareto optimal MM, the blood’s optimal residence time within the capillary is roughly 1*/*2 of the cycle time: *f*_0_*l/U* = (*π/*4)^3^≈ 0.48. We use the kinematic metabolic rate (Eq. 6) when deriving the energetic metabolism of ‘large’ organisms by adding up the MMs so that they are fuelled in one organism and work together.

## 4 Adding up Metabolic Modules

As Figure 3 reveals, the general metabolic theory has the total number *n* of MMs as the most relevant coulter parameter with *n* = 1 for protists and planarians, *n* = 10^3^ for ectotherms, and *n* = 10^8^ for endotherms including mammals. As shown by Figure 2, the general theory based on one MM (*n* = 1) covers the *diffusion asymptote* of metabolism valid for protists and also the *convection asymptote* valid for planarians. All organisms share the same *diffusion asymptote* but differ in the *convection asymptote*.

Physiological data reported in literature give the parameter for the general metabolic theory and the results plotted as black solid lines in Figure 3. Metabolism happens in cells. This repeated reminder is needed when applying and parameterizing the theory derived above. Since cells are the basic building blocks of all organisms, the measures *ϱ, e, k, 𝒟* must be equal at least in the order of magnitude for all organisms. The reaction rate of glucose oxidation is *k* 0.1≈ s^−1^ [34]. With the known reaction rate, the measured metabolic rate of protists [35], [36] (Fig. 3), the mass density *ϱ* = 1000 kg*/*m^3^, and the *diffusion asymptote* (Eq. 1), we derive the energy density to be *e* = |Δ*G*^*°*^| Δ*c* ≈ 57.4 kJ*/*m^3^. The change in Gibbs’ energy for glucose oxidation is Δ*G*^*°*^ = −2.87 MJ mol^−1^ [37]. Therefore, the oxygen concentration is Δ*c* = 0.02 mol*/*m^3^.

For the MM the dispersion coefficient *D* and the critical shear rate 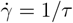 are physiological constants. To prevent primary haemostasis, the required shear rate within the capillaries of humans is 10^2^ … 10^3^ 1*/*s [29], [38]. In the absence of further data, we assume *τ*≈ 1 ms for all organisms, recognising only a weak sensitivity in the general metabolic theory and the associated asymptotes (Eqs. (2) to (10)), especially in comparison to the sensitivity to other physiological constants such as *k* and *D*. I am aware that the flow in the gastric cavity of planarians differs significantly from the blood flow in the capillaries of humans. The molecular coefficient of diffusion in dilute solutions is 𝒟=10^−9^ m^2^*/*s [39]. Calibrating the general theory for *n* = 1 to the data of protists [35], [36] and planarians [7] yields *D* = 10^−3^𝒟. As expected, this dispersion coefficient is smaller than the molecular diffusion coefficient of molecules in dilute solution.

Evolution has overcome the growth constraint in planarians through one MM that is up-scaled. As cells are up-numbered MMs are up-numbered. Nature allows further growth by multiplying MMs in an organism. In this up-numbering, *n≫* 1 MMs are fuelled in one organism and work together. Using the above parameters and the experimental data for ectotherms and endotherms, the total numbers *n* = 10^3^ and *n* = 10^8^ were identified, which predict the basal energetic metabolic rate, as shown in Figure 3 with black solid lines. The general theory covers the complete mass ranges from one picogram to 100 tons.

Up-numbering gives the transformation from volume to mass, *V*→ *m*, and from kinematic metabolic rate to energetic metabolic rate, *Q*→ *Ė*: *m* = *ϱnV, Ė* = *a ne Q*. The activity factor *a* is introduced at the same time. This indicates the ratio of active to basal volume flow. As a first approximation, this ratio is equal to the ratio of heart rates. The dashed solid line in Figure 3 shows the predicted metabolic rate of active mammals fitting measured data [35], [40] including that of elite athletes rowing a 2000 m Olympic regatta [41].

## 5 Kleiber’s Law as Special Case of the General Theory

Kleiber’s law, i.e., the scaling of the metabolic rate, the scaling of the heart rate of mammals, and the scaling of the life span of mammals is given in the following. Kleiber’s law, *Ė* ∝ *m*^3*/*4^, is gained from the *convection asymptote* (Eq. 6) by up-numbering:

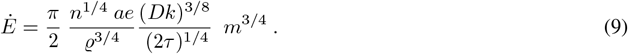

The *convection asymptote* (Eq. 9) is shown as a dotted line in Figure 3 for mammals with *a* = 1. The scaling of the basal heart rate with body mass *f* ∝ *m*^−1*/*4^ is derived as follows. The dimensionless product *f V/Q* = *A* = const. must be scale-invariant as discussed above due to Bridgman’s postulate [8]. With this and Eq. (8) the heart frequency is derived as

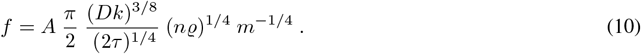

If one assumes that an organism bears only a limited total number Ω of blood circulations during the life cycle from birth to death, then the life span *T* is given by multiples of the residence time *T* = Ω *l/U*. Experimental data for mammals, cf. Fig. 5, show that *f V/Q* = *A* ≈ 10^−2^ and the total number of cycles Ω≈ 1.51 * 10^11^. Hence, it follows that the total number of heart beats *N* = *f T* during a lifetime are the same for mouse and elephant: *N* = (*π/*4)^3^ *A*Ω. This result of the general theory of metabolism is confirmed by experience [10]. Each mammal has a budget of *N* = 0.48 *A*Ω = (0.73 *±* 0.56) * 10^9^ heart beats.

## 6 Outlook

The paper contributes to the perception of evolution and metabolic scaling. The general theory of metabolism for a given organism’s size and taxonomy is quantitative in contrast to known theories and is consistent with cell metabolism. Despite the model’s simplicity, the metabolic rate prediction is surprisingly accurate at least on the logarithmic scale ranging from one picogram to hundreds of tons. The general theory of metabolism covers two asymptotes, the *diffusion asymptote* and the *convection asymptote*. The latter yields Kleiber’s laws. All physicochemical and physiological model parameters are in the expected range.

The simplicity of the model is reflected in the few assumptions that are necessary for the theory. The simplicity is therefore an advantage over existing models. The questions remaining open motivate further research. One such question is whether all MMs are supplied in parallel or whether a portion is also supplied in series. If the latter were the case, the volume flow would be reduced. In fact, this would be consistent with the relative heart volume.

## References

[1] Kleiber, M. Body size and metabolism. Hilgardia 6(11), 315–353 (1932). DOI: 10.3733/hilg.v06n11p315

[2] Galilei, G. and Elzevier. Discorsi e dimostrazioni matematiche, intorno à due nuoue scienze (1638)

[3] Smil, V. Laying down the law. Nature 403(6770), 597–597 (2000). DOI: 10.1038/35001159

[4] Murray, C. D. The physiological principle of minimum work: I. the vascular system and the cost of blood volume. Proceedings of the National Academy of Sciences 12(3), 207–214 (1926). DOI: 10.1073/pnas.12.3.207

[5] Taber, L. A., Ng, S., Quesnel, A. M., Whatman, J., and Carmen, C. J. Investigating Murray’s law in the chick embryo. Journal of Biomechanics 34(1), 121–124 (2001). DOI: 10.1016/S0021-9290(00)00173-1

[6] Painter, P. R., Edén, P., and Bengtsson, H.-U. Pulsatile blood flow, shear force, energy dissipation and Murray’s Law. Theoretical Biology and Medical Modelling 3(1), 31 (2006). DOI: 10.1186/1742-4682-3-31

[7] Thommen, A., Werner, S., Frank, O., Philipp, J., Knittelfelder, O., Quek, Y., Fahmy, K., Shevchenko, A., Friedrich, B. M., Jülicher, F., and Rink, J. C. Body size-dependent energy storage causes Kleiber’s law scaling of the metabolic rate in planarians. eLife 8, e38187 (2019). DOI: 10.7554/eLife.38187

[8] Bridgman, P. W. Dimensional analysis (Yale University Press 1922). urn:oclc:record:1042888750

[9] Schmidt-Nielsen, K. Animal physiology, 5th ed (Cambridge University Press 2008). ISBN: 978-0-521-57098-5

[10] Levine, H. J. Rest heart rate and life expectancy. Journal of the American College of Cardiology 30(4), 1104–1106 (1997). DOI: 10.1016/S0735-1097(97)00246-5

[11] Ballesteros, F. J., Martinez, V. J., Luque, B., Lacasa, L., Valor, E., and Moya, A. On the thermodynamic origin of metabolic scaling. Nature Scientific Reports 8(1), 1448 (2018). DOI: 10.1038/s41598-018-19853-6

[12] West, G. B., Brown, J. H., and Enquist, B. J. A general model for the origin of allometric scaling laws in biology. Science 276(5309), 122–126 (1997). DOI: 10.1126/science.276.5309.122

[13] Banavar, J. R., Maritan, A., and Rinaldo, A. Size and form in efficient transportation networks. Nature 399(6732), 130–132 (1999). DOI: 10.1038/20144

[14] Savage, V. M., Deeds, E. J., and Fontana, W. Sizing up allometric scaling theory. PLoS Computational Biology 4(9), e1000171 (2008). DOI: 10.1371/journal.pcbi.1000171

[15] Hertz, H. Die Prinzipien der Mechanik (Johann Ambrosius Barth, Leipzig 1894)

[16] Christiansen, F. V. Heinrich Hertz’s neo-Kantian philosophy of science, and its development by Harald Høffding. Journal for General Philosophy of Science 37(1), 1–20 (2006). DOI: 10.1007/s10838-006-0486-0

[17] Paqué, R. Das Pariser Nominalistenstatut (Walter de Gruyter, Berlin 1970). DOI: 10.1515/9783110816785

[18] Dawson, T. H. Scaling laws for capillary vessels of mammals at rest and in exercise. Proceedings of the Royal Society of London. Series B: Biological Sciences 270(1516), 755–763 (2003). DOI: 10.1098/rspb.2002.2304

[19] Schmalhausen, I. I. Evolution and cybernetics. Evolution 14(4), 509–524 (1960). DOI: 10.1111/j.1558-5646.1960.tb03117.x

[20] Cavagna, G. A. and Kaneko, M. Mechanical work and efficiency in level walking and running. The Journal of Physiology 268(2), 467–481 (1977). DOI: 10.1113/jphysiol.1977.sp011866

[21] Lighthill, M. J. Note on the swimming of slender fish. Journal of Fluid Mechanics 9(2), 305–317 (1960). DOI:10.1017/S0022112060001110

[22] Lighthill, J. Aerodynamic aspects of animal flight. In: Wu, T. Y.-T., Brokaw, C. J., and Brennen, C. (eds), Swimming and Flying in Nature (Springer US, Boston, MA 1975). 423–491. DOI: 10.1007/978-1-4757-1326-8_1

[23] Glazier, D. S. Variable metabolic scaling breaks the law: from ‘Newtonian’ to ‘Darwinian’ approaches. Proceedings of the Royal Society B: Biological Sciences 289(1985), 20221605 (2022). DOI: 10.1098/rspb.2022.1605

[24] Goethe, J. W. v. Faust (Cotta’sche Verlagsbuchhandlung, Tübingen 1808)

[25] Pelz, P. F., Groche, P., Pfetsch, M. E., and Schaeffner, M. Mastering uncertainty in mechanical engineering (Springer 2021). DOI: 10.1007/978-3-030-78354-9

[26] Rams, D. Less but better, 6th ed (Die Gestalten Verlag 2016). ISBN: 978-3-89955-525-7

[27] Saint-Exupéry, A. d. Terre des Hommes (Gallimard 1939). ISBN: 978-2-07-036021-5

[28] Kolmogorov, A. N. Local structure of turbulence in an incompressible viscous fluid at very high Reynolds numbers. Doklady Akademii Nauk SSSR 31, (1941)

[29] Gogia, S. and Neelamegham, S. Role of fluid shear stress in regulating VWF structure, function and related blood disorders. Biorheology 52(5-6), (2015). DOI: 10.3233/BIR-15061

[30] Khamassi, J., Bierwisch, C., and Pelz, P. Geometry optimization of branchings in vascular networks. Physical Review E 93(6), 062408 (2016). DOI: 10.1103/PhysRevE.93.062408

[31] Mazzag, B. M., Tamaresis, J. S., and Barakat, A. I. A model for shear stress sensing and transmission in vascular endothelial cells. Biophysical Journal 84(6), 4087–4101 (2003). DOI: 10.1016/S0006-3495(03)75134-0

[32] Barakat, A. I., Lieu, D. K., and Gojova, A. Secrets of the code: do vascular endothelial cells use ion channels to decipher complex flow signals? Biomaterials 27(5), 671–678 (2006). DOI: 10.1016/j.biomaterials.2005.07.036

[33] Taylor, G. I. Dispersion of soluble matter in solvent flowing slowly through a tube. Proceedings of the Royal Society of London. Series A. Mathematical and Physical Sciences 219(1137), 186–203 (1953). DOI: 10.1098/rspa.1953.0139

[34] Tao, Z., Raffel, R. A., Souid, A.-K., and Goodisman, J. Kinetic studies on enzyme-catalyzed reactions: oxidation of glucose, decomposition of hydrogen peroxide and their combination. Biophysical Journal 96(7), 2977–2988 (2009). DOI: 10.1016/j.bpj.2008.11.071

[35] Hemmingsen, A. M. Energy metabolism as related to body size and respiratory surfaces, and its evolution. Reports of the Steno Memorial Hospital and then Nordisk Insulinlaboratorium Gentofte, Denmark 9 (1960)

[36] DeLong, J. P., Okie, J. G., Moses, M. E., Sibly, R. M., and Brown, J. H. Shifts in metabolic scaling, production, and efficiency across major evolutionary transitions of life. Proceedings of the National Academy of Sciences 107(29), 12941–12945 (2010). DOI: 10.1073/pnas.1007783107

[37] Tran, Q. H. and Unden, G. Changes in the proton potential and the cellular energetics of Escherichia coli during growth by aerobic and anaerobic respiration or by fermentation. European Journal of Biochemistry 251(1), 538–543 (1998). DOI: 10.1046/j.1432-1327.1998.2510538.x

[38] Khamassi, J. Untersuchung der Adhäsion von Thrombozyten mittels Partikelsimulation und mikrofluidischer Experimente (Shaker Verlag 2018). ISBN: 978-3-8440-6325-7

[39] Probstein, R. F. Physicochemical hydrodynamics (Butterworth-Heinemann 1989). ISBN: 978-0-7506-9401-8

[40] West, G. Complexity, regularity, diversity and scale of diversity (Santa Fe Institute, San Francisco 2018). URL: https://www.youtube.com/watch?v=WcwqcKkkkcg

[41] Pelz, P. F. and Vergé, A. Validated biomechanical model for efficiency and speed of rowing. Journal of Biomechanics 47(13), 3415–3422 (2014). DOI: 10.1016/j.jbiomech.2014.06.037

[42] Witting, L. Inevitable evolution: back to the origin and beyond the 20th century paradigm of contingent evolution by historical natural selection. Biological Reviews 83(3), 259–294 (2008). DOI: 10.1111/j.1469-185X.2008.00043.x

[43] Seymour, R. S. and Blaylock, A. J. The principle of laplace and scaling of ventricular wall stress and blood pressure in mammals and birds. Physiological and Biochemical Zoology 73(4), 389–405 (2000). DOI: 10.1086/317741

